## Supplemental for "Electrical Readout Strategies of GFET Biosensors for Real-World Requirements"

Evaluation of Electrical Readout Strategies for GFET-Based Viral Biosensing

**Figure S1** – Fresh PDMS vs. Aged PDMS vs. Acrylic well initial Dirac point voltages. Acrylic wells show a lower initial Dirac point with less variability across numerous devices.

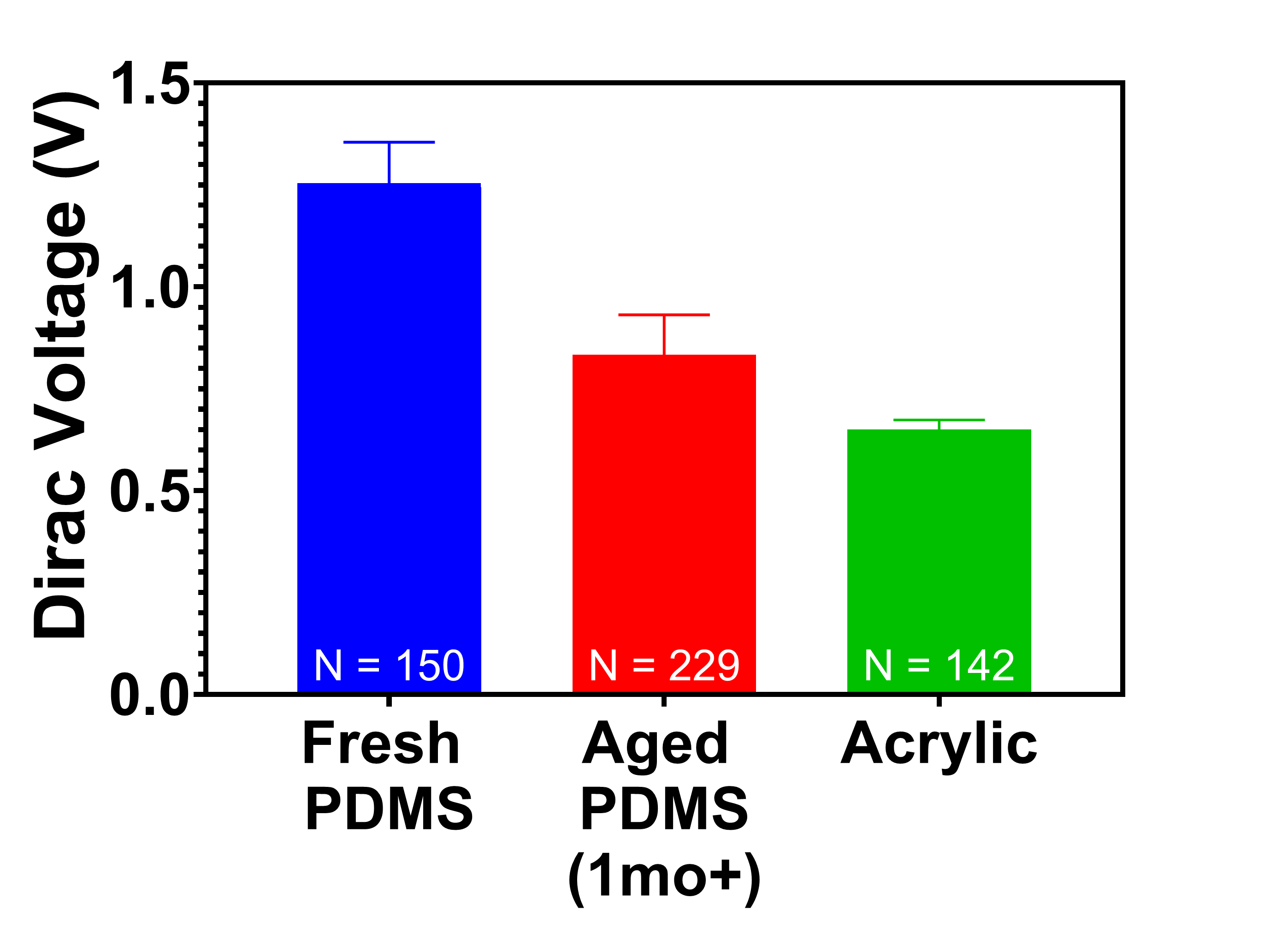

**Figure S2** – Clover well design. a) placed well with no liquid. Notice small cross in center of well fabricated on-chip as a marker. b) Same will with each leaf containing different colored dyed PBS. Notice cross still visible showing liquids not entering the center of the well and mixing with each other. c) liquid added to center of well above cross spreads to all clover leaves.

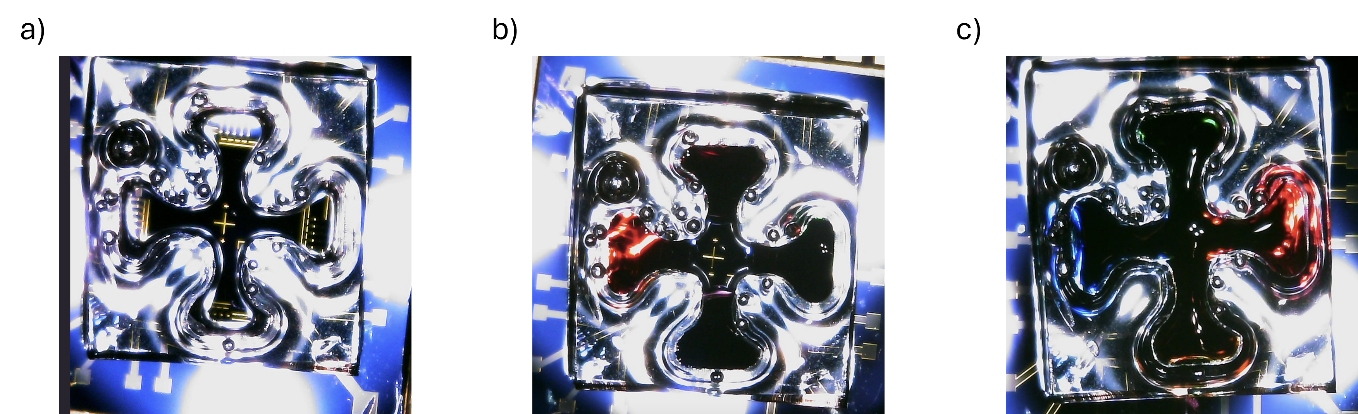

**S3** – Aptamer Linking process

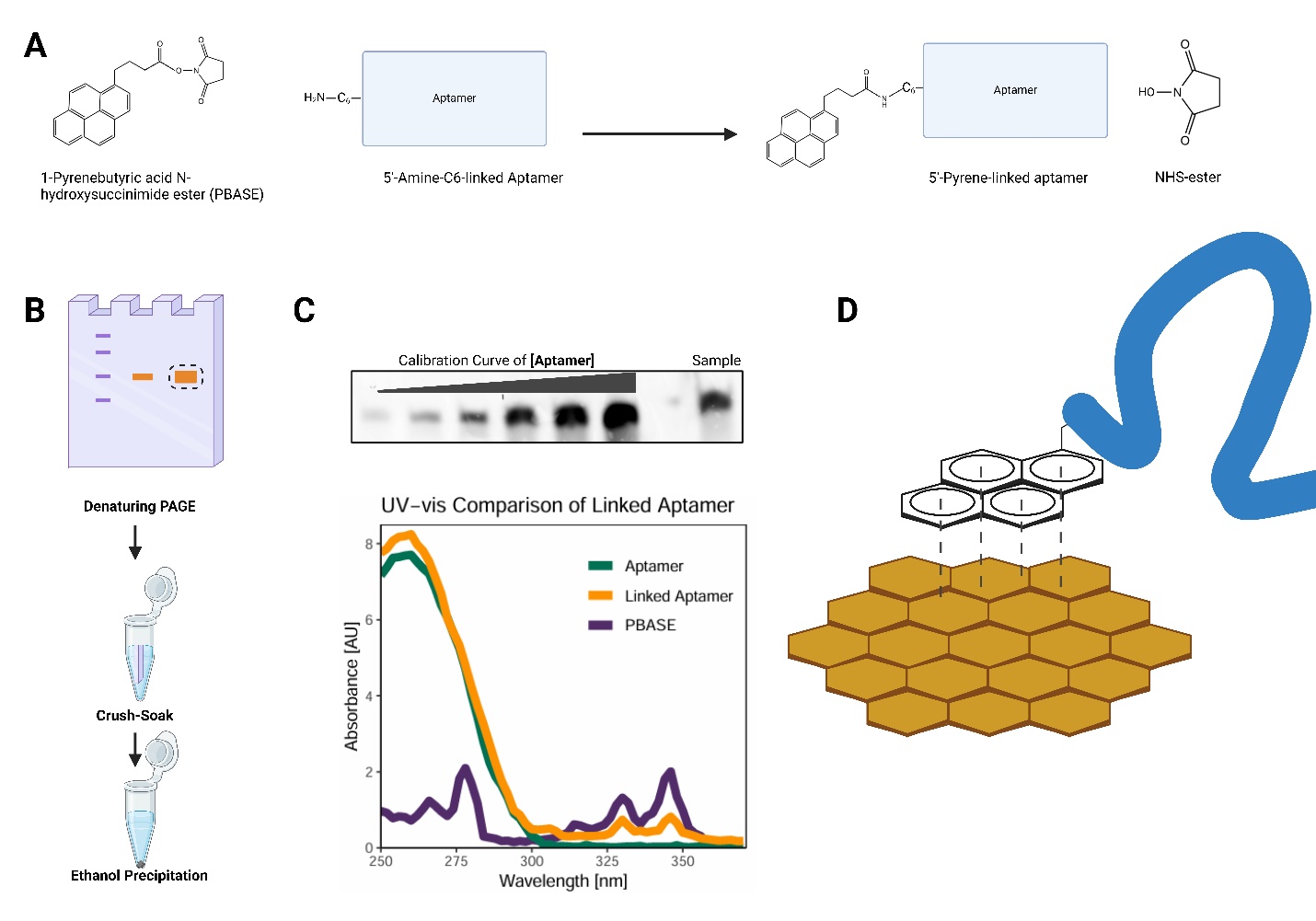

**Figure S3- Scheme of Pyrene-Aptamer conjugation reaction.**

**A,** 1-Pyrenebutyric acid N-hydroxysuccinimide (PBASE) conjugation to an aptamer with a 5’ C_6_-amine modification. In slightly alkaline conditions, the NHS ester reacts with the primary amine, linking the pyrene to the aptamer and releasing NHS. **B,** To purify pyrene-linked aptamer, the reaction is separated via denaturing PAGE and stained with ethidium bromide to visualize DNA using transillumination (365 nm). Linked aptamer is excised and extracted from gel via diffusion and subsequently ethanol precipitated. **C,** Quality checks for pyrene-linked aptamer include gel densitometry to quantify linked aptamer via a calibration curve (5, 10, 25, 50, 75, 150 pmol of unlinked aptamer visualized ethidium bromide) and UV-vis spectra to confirm PBASE conjugation. The UV-vis spectra of 10 pmol PBASE, 10 pmol linked aptamer, and 10 pmol aptamer are compared. **D,** Functionalization of graphene. The linked aptamer is then used for concentration-dependent assays of specified targets. Figure created with BioRender.com

**Methods:**

Conjugation

The *in vitro* pyrene-aptamer conjugation differs from the previous linking reaction^1^ in that it was performed in a microcentrifuge tube instead of on the graphene surface. The goal of this acyl transfer is for the NHS ester group of the 1-Pyrenebutyric acid N-hydroxysuccinimide (PBASE) to interact with the amine group attached to the 5’ end of the aptamer (**Figure S3A**), resulting in a pyrene-linked aptamer that can be attached to the graphene. To achieve this, we adapted a manufacturer’s protocol (<https://www.sigmaaldrich.com/US/en/technical-documents/protocol/genomics/pcr/nhs-ester-oligonucleotide-conjugation>) as described previously (Geiwitz et al. 2024)

Gel Extraction and Ethanol Precipitation

After conjugation reaction, samples were separated via a 6 % polyacrylamide 7.5 M urea gel (National Diagnostics, EC-830) at a constant 35 mA for 90 minutes (**Figure S3B**). Linked RNA was visualized via post-staining with ethidium bromide and imaged at 365 nm. Linked nucleic acid fragments were excised using a razor and gel slices were directly transferred into a crush soak solution (5 M NaCl, pH 7.5, 1 M Tris·HCl pH 7.5, 0.5 M EDTA pH 8), and pyrene-linked aptamers were extracted via diffusion (5:1 ratio of crush soak buffer volume: gel fragment mass incubated on a rocker for an hour at 37°C). To remove any small fragments of acrylamide resulting from the excision and diffusion process, we pre-precipitated by adding 20% ethanol by volume, followed by centrifugation at 13,200 rpm for 15 minutes. Subsequently, 500 μL of the supernatant was transferred into a tube of 1 mL of ice-cold ethanol and 1 μL of glycogen to enable ethanol precipitation of the linked nucleic acid. Complete extraction of nucleic acids from the gel was ensured via repeating diffusion and precipitation procedures up to three times for the same gel piece. Linked aptamers in ethanol were incubated at -20°C before centrifugation at 13,200 rpm for 15 minutes to pellet precipitated linked aptamer. The supernatant was removed, and the pellet was allowed to dry before reconstitution in H_2_O.

Gel Densitometry

The pyrene group of the PBASE interferes with characteristic ssDNA absorbance at 260 nm^2^. Therefore, concentration was obtained through quantitative gel densitometry (**Figure S3C**). Using a 6 % polyacrylamide 7.5 M urea gel, a standard curve of known aptamer concentrations and an aliquot of the linked aptamer underwent electrophoresis, ethidium bromide staining, and quantitative imaging [General Electric Typhoon FLA 9500]. A calibration plot of the standard curve band density (raw volume) vs. concentration was used to calculate the concentration of the linked aptamer. The pyrene-conjugated aptamer was then aliquoted to 50 μL of 10 μM for downstream applications.

UV-vis Spectrophotometry

PBASE has characteristic UV adsorption peaks at 325 nm and 341 nm^3^ absent in aptamers. Therefore, comparing UV-vis peaks from PBASE, the unlinked aptamer, and a sample of linked aptamer ensures a successful linking reaction (**Figure S3C**).

**Figure S4** – Dirac point shift as a function of gate voltage transfer curves for Influenza A hemagglutinin; a) with applied gate voltage at max transconductance point during target incubation and b) with no applied field during target incubation. With applied field, there were no significant shifts in Dirac point voltage until a much higher than shifts were observed with no field applied.

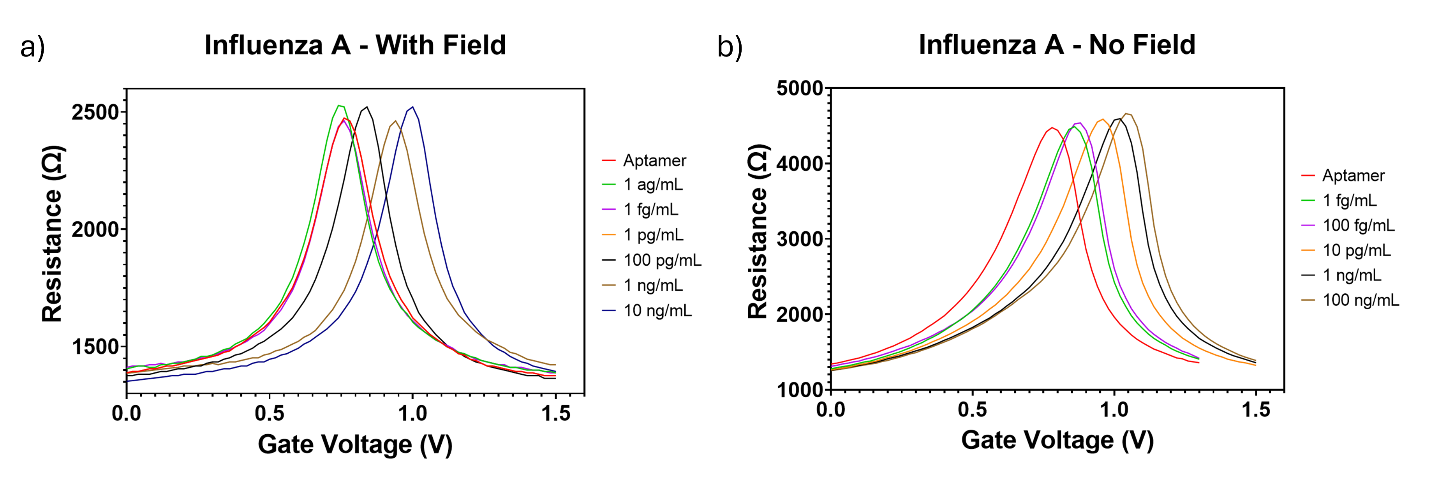

1.	Kumar, N. *et al.* Rapid, Multianalyte Detection of Opioid Metabolites in Wastewater. *ACS Nano* 16, 3704–3714 (2022).

2.	Maeda, H., Inoue, Y., Ishida, H. & Mizuno, K. UV Absorption and Fluorescence Properties of Pyrene Derivatives Having Trimethylsilyl, Trimethylgermyl, and Trimethylstannyl Groups. *Chem. Lett.* **30**, 1224–1225 (2001).

3.	Bao, Q. *et al.* Graphene–Polymer Nanofiber Membrane for Ultrafast Photonics. *Adv. Funct. Mater.* **20**, 782–791 (2010).
